# DynOmics to identify delays and co-expression patterns across time course experiments

**DOI:** 10.1101/076257

**Authors:** Jasmin Straube, Bevan Emma Huang, Kim-Anh Lê Cao

## Abstract

Dynamic changes in biological systems can be captured by measuring molecular expression from different levels (*e.g.*, genes and proteins) across time. Integration of such data aims to identify molecules that show similar expression changes over time; such molecules may be co-regulated and thus involved in similar biological processes. Combining data sources presents a systematic approach to study molecular behaviour. It can compensate for missing data in one source, and can reduce false positives when multiple sources highlight the same pathways. However, integrative approaches must accommodate the challenges inherent in ‘omics’ data, including high-dimensionality, noise, and timing differences in expression. As current methods for identification of co-expression cannot cope with this level of complexity, we developed a novel algorithm called DynOmics. DynOmics is based on the fast Fourier transform, from which the difference in expression initiation between trajectories can be estimated. This delay can then be used to realign the trajectories and identify those which show a high degree of correlation. Through extensive simulations, we demonstrate that DynOmics is efficient and accurate compared to existing approaches. We consider two case studies highlighting its application, identifying regulatory relationships across ‘omics’ data within an organism and for comparative gene expression analysis across organisms.

## Introduction

High-throughput ‘omics’ platforms such as transcriptomics, proteomics, and metabolomics enable the simultaneous monitoring of thousands of biological molecules (transcripts, proteins, and metabolites), typically through a single static experiment.^1^ The recent decrease in cost of such technological platforms has made possible the study of dynamic biological processes by instead quantify molecules at several time points. This allows deeper insight into the behaviour of the molecules in situations ranging from developmental processes to drug response. These time course ‘omics’ experiments enable the identification of regulators, and may give a better understanding of the structure and dynamics of biological systems.

The statistical analysis of dynamic ‘omics’ experiments is difficult. Applying traditional statistical methods for static experiments is limited, since each time point will be treated as independent, ignoring potentially important correlations between sampling times. Indeed, realising the potential power offered in time course studies to investigate a wide variety of changes is nontrivial. Analytical challenges are further complicated by noise, small sample sizes per time point, and few sampled time points. In the past decade, several methods have been proposed to analyse time course ‘omics’ data, with a particular focus on microarray and RNA-Seq data. These methods perform differential expression analysis using spline fitting,^2, 3^ Bayesian methods,^4–7^ Gaussian processes,^8–10^ and a two-step regression approach (maSigPro^11^). Other methods focus on clustering expression profiles to identify co-expressed trajectories, *e.g.*, a subset of molecules for which expression changes occur simultaneously across time.^3, 7, 12–16^ Targeted co-expression analysis can also be performed using various model-based applications to retrieve data sets from databases given specific query data.^17–19^ Finally, a third category of methods was proposed based on biological pathway analysis^20, 21^ see the detailed review of Spies *et al.*^22^ Co-expression analysis can provide valuable insight into the role of molecules during biological processes,^23–25^ but faces significant challenges in dealing with different types of ‘omics’ and their variation in molecular response times. These timing differences or delays in the initiation or suppression of molecule expression are a common phenomenon in biology and occur across both different molecular levels and organisms. For example, the study of regulatory processes after environmental changes has revealed that there is often a measurable delay from the time of signal introduction to molecular response.^23, 24, 26–28^ This can result from differences in the reaction kinetics between an enzyme and its substrate, presence of an inhibitor, or altered binding affinities of transcription factors. Such processes can be studied through time course miRNA and mRNA data, since miRNAs play an important role in gene translation regulation in many organisms, through either mRNA translation inhibition or mRNA degradation.^29^ This ability of miRNAs to fine tune gene expression and translation in a broad range of important biological processes is of broad interest in medicine.^30–32^ While correlation analysis is frequently used to analyse miRNA-mRNA time course data,^33, 34^ it may have limited power in situations where delayed dynamic expression changes of miRNAs relative to mRNA have been observed.^30, 35^

Delays can also hinder gene expression comparisons across organisms, since even highly conserved processes may vary in timing. The pre-implantation embryonic development (PED) is a highly conserved process across mammals, reflected through the progression of the same morphologic stages.^36^ Nevertheless, attempts to compare PED in mammals based on gene expression data have faced challenges due to differences in timing of genome activation and regulatory processes.^37^ Hence ignoring these delays in co-expression analysis can mask true associations; the first step should instead be to detect and quantify the time delay between molecules. This will enable identification of functionally related molecules regardless of the differences in the timing of expression changes, as well as allowing quantification of similarities and differences between the observed responses in more detail.

To date, very few methods for time course ‘omics’ data account for time delay between molecule expression levels. Aijo *et al.*^10^ recently proposed DynB, a set of methods based on Gaussian processes, to quantify RNA-Seq gene expression dynamics. This allows rescaling of time profiles, but only between replicates (*i.e.*, at the sample level) rather than at the molecule expression level. The most commonly used approach for molecules is to consider the Pearson correlation,^34, 38^ despite its obvious limitations for detecting co-expressed molecules when their expression change occurs at different time points. Lagged Pearson correlation, *a.k.a.* Pearson cross-correlation for lagged time series, circumvents this limitation by introducing artificial delays or lags in the time expression profiles for every possible time shift. The method eventually applies the delay that maximises the correlation with the original profile, but can be prone to overestimation of delay.

More sophisticated approaches for time course ‘omics’ data come at the expense of computational cost. Shi *et al.*^39^ proposed an probabilistic model based on multiple datasets tabular combinations to identify pairwise transcription factor and gene (TF-G) pairs under different experimental conditions. This approach has shown to reduce false positive predictions but requires a time consuming learning step on existing and known TF-G pair data.^39^ Dynamic Time Warping (DTW)^40–42^ is an algorithm that aligns the time points of two trajectories to minimise the distance between them. It can therefore identify similarities between trajectories which may vary in phase and speed. One variation, DTW4Omics^24^ identifies co-expressed molecules with a permutation test, but this can be computationally expensive. An alternate approach^25^ utilises a combined statistic based on Hidden Markov Models (HMM) and Pearson correlation. HMMs are trained on a set of trajectories where a distribution of values is considered for each time point. This generates a probability to observe a trajectory under the trained model that can tolerate small delays. While promising, this approach cannot detect large delays. Additionally, both it and DTW4Omics can only identify positively correlated trajectories, requiring heavier computational costs to exhaustively explore potential associations.

While integrating time course experiments from different ‘omics’ functional levels is the key to identifying dynamic molecular interactions, its challenging nature has thus far prevented much methodological development. Difficulties lie not only in the computation required by complex algorithms, but also in variation in types of correlation, levels of noise in the expression profiles, and the delays themselves.

We present DynOmics, a novel algorithm to detect, estimate, and account for delays between ‘omics’ time expression profiles. The algorithm is based on the fast Fourier transform (FFT),^43^ which has already been shown to successfully detect periodically expressed genes in transcriptomics experiments.^44–46^ By combining the FFT angular difference between reference and query trajectories with lagged Pearson correlation, we are able to characterise the direction and magnitude of delay, whether the reference and query are positively or negatively correlated. After accounting for the estimated delays, similar profiles can be clustered for further insight. Simulation results show that DynOmics outperforms current methods to detect time shift, both in terms of sensitivity and specificity. We apply it to two biological case studies: one focusing on the integration of miRNAs and mRNAs in mouse lung development, and one on the conservation of gene molecular processes across multiple organisms (mouse, bovine, human) during PED. In both cases, DynOmics is able to unravel timing differences between ‘omics’ functional levels, demonstrating its wide applicability. DynOmics is available as an R package on bitbucket.^47^

## Material and Methods

The expression changes of molecules monitored in time course experiments often form simple temporary, sustainable or cyclic patterns that can be modelled as mixtures of oscillating/cyclic patterns using the discrete Fourier transform (FT).^48^ We introduce DynOmics, a novel method that first converts trajectories to the frequency domain using the FFT, from which it extracts the frequency of the main cyclic pattern. Condensing the trajectory to information on the main frequency is then used to identify whether two trajectories are related or *associated*, while ignoring the noise in each time expression profile.

### Fourier Transform

For a given time series *x* = (*x*_1_,…,*x*_*t*_,…*x*_*T*_), measured at time points *t* = 1,…*T*, the FT decomposes *x* into circular components or cyclic patterns for each frequency *k* = 1, …, *T* − 1 as:

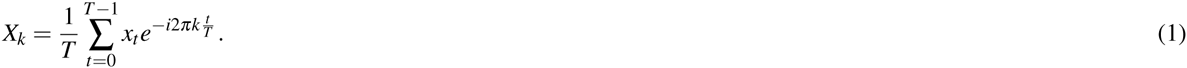

As the amplitude at frequency *k* = 0 simply describes the y-axis offset (*i.e.*, the global differences of expression levels), this frequency is not included in our analysis context. Equation (1) can be written with polar coordinates with real part *a* and imaginary part *b* as *X*_*k*_ = *a*_*k*_ + *b*_*k*_*i*.

For each frequency *k* = 1, …, *T* − 1 we can calculate the amplitude *r*_*k*_ of the component as 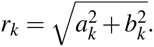 The amplitude reflects the contribution of the *k*^*th*^ cyclic pattern to the overall trajectory, and the pattern with maximum amplitude 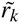 describes the main shape of the time series. The argument *Arg*(*X*_*k*_) is the offset of the cyclic pattern, defined as:

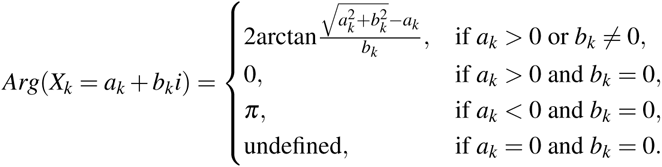

We can transform the argument to the phase angle (delay) *φ*_*k*_ in degrees by:

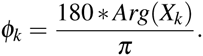

Together, the amplitude and phase angle describe each frequency component, and the set of these quantities is known as the frequency domain representation.

### DynOmics

We describe DynOmics, a novel method to estimate delays between a reference *x* and query *y* given in frequency domain representation. First, we identify *K* as the frequency of the pattern with maximum amplitude for *x*, *i.e.*, the main reference pattern frequency. Then, for both *x* and *y*, we extract phase angles at this frequency *ϕ*^*x*^ = *ϕ*_*xK*_,*ϕ*^*y*^ = *ϕ*_*yK*_ and define Δ_*xy*_ = *ϕ*^*x*^ − *ϕ*^*y*^ as the difference between the phase angles. In FFT literature, ∆_*xy*_ is often expressed in the range of [−180,180]. To simplify representation in DynOmics, when Δ_*xy*_ < 0, we add 360 so that Δ_*xy*_ is in the range of 0 to 359. Δ_*xy*_ indicates both the sign of the correlation between *x* and *y* and the sign of the delay, as seen in Figure 1. The trajectories *x* and *y* can be either positively (Figure 1 **abf**) or negatively correlated (Figure 1 **cde**), with a delay that we refer to as *negative*, *i.e.*, the reference *x* is prior to the query *y* (Figure 1 **be**) or *positive*, *i.e.*, the reference *x* is delayed with respect to the query *y* (Figure 1 **cf**). Specific angular difference cases include when ∆_*xy*_ = 0 (positive correlation, but no delay, Figure 1 **a**) and when ∆_*xy*_ = 180 (negative correlation, no delay, Figure 1 **d**). We can estimate the delay between two trajectories based on the FT frequency, the length of the time series and ∆_*xy*_ as

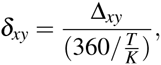

**Figure 1.**
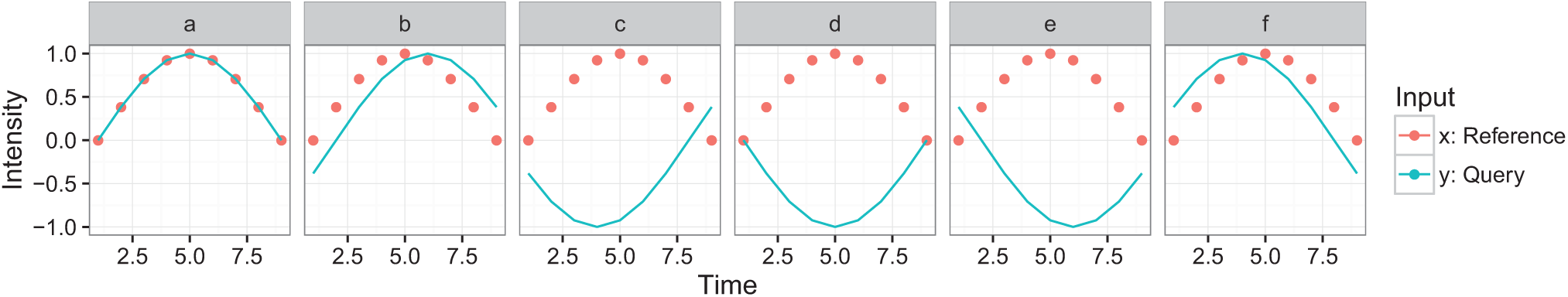
Relationship between angular differences, correlation and delay for a reference trajectory *x* (red dots) and a query trajectory *y* (green line). The trajectories are **a**) positively correlated with no delay (∆_*xy*_ = 0); **b**) positively correlated with negative delay (0 *<* ∆_*xy*_ 90); **c**) negatively correlated with positive delay (90 *<* ∆_*xy*_ < 180); **d**) negatively correlated with no delay (∆_*xy*_ = 180); **e**) negatively correlated with negative delay (180 *<* ∆_*xy*_ < 270); **f**) positively correlated with positive delay (270 *≤* ∆_*xy*_ < 360).

where *δ*_*xy*_ ranges from 0 to 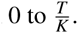. In order to keep delay estimates and hence signal shifts as small as possible, we collapse these values to the range of 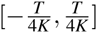 by setting 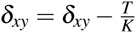 when 270 *≤* ∆ _*xy*_ *≤* 359, and 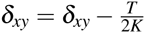 when 90 *≤* ∆_*xy*_ *≤* 270. We note that for query profiles either positively or negatively correlated with the reference, this means that *δ*_*xy*_ < 0 represents positive delay, and *δ*_*xy*_ > 0 represents negative delay.

### Using lagged Pearson correlation to increase accuracy in delay estimation

The delay estimate based on the angular difference presented above is based on approximating both the reference and query by the pattern at the main frequency for the reference. This approximation works well when the signals from both query and reference are dominated by that main pattern and relatively ‘noise-free’. However, when multiple frequencies have substantial contributions to the overall signal, we can improve the estimate by maximising the lagged Pearson correlation coefficient over small perturbations in the delay.

Specifically, let *δ*_0_ = ⌊*δ*_*xy*_⌋ denote our initial delay estimate, rounded to the closest integer. Let *ℒ* be a set of lags {*l*} representing perturbations to this initial delay estimate. For each lag, we construct trajectories *x*_*l*_ and *y*_*l*_ by shifting the original trajectories so that if *l* < 0,*x*_*l*_ = *x*_1+|*l*|_,…,*x*_*T*_ and *y*_*l*_ = *y*_1_,…,*y*_*T*−|*l*|_; if *l* > 0, conversely, *x*_*l*_= *x*_1_,…,*x*_*T*−|*l*|_ and *y*_*l*_ = *y*_1+|*l*|_,…,*y*_*T*_. The (lagged) Pearson correlation coefficient between the two trajectories *x*_*l*_ and *y*_*l*_ is defined as:

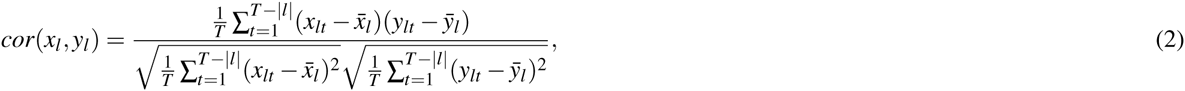

where 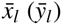 is the sample mean across time points for each trajectory. We determine the optimal delay for a given set of lags as that for which the Pearson correlation coefficient is maximised:

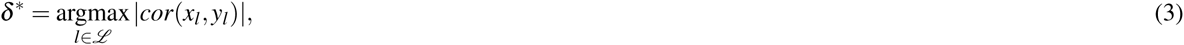

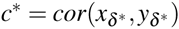 then used to assess the strength and direction of the association.

We consider two sets *ℒ*_1_, *ℒ*_2_ of lags which represent perturbations of *δ*_0_:

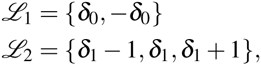

where *δ*_1_ is the result of the optimization over *l* Ϻ *ℒ*_1_. These two optimisations thus allow us to compare the initial estimate with that in the opposite direction, and then with delays in a local neighbourhood. While the optimisations do increase the computation required, our restriction to local perturbations minimises the additional computation while improving the estimate in the presence of noise.

### Sensitivity and specificity of DynOmics compared to other studies

We compared DynOmics performance with current available methods (DTW4Omics,^24^ Pearson and lagged Pearson correlation) using measures of sensitivity and specificity while identifying associations in simulated data. The simulated data were generated based on similar scenarios to^25^ with different parameters. We simulated different expression patterns with different delays (−2, 1, 0, 1, 2) as well as different noise levels 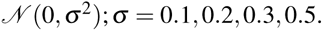 We also simulated different number of time points (7 and 14 times points). A number of 7 times points characterises best conventional ‘omics’ time course experiments.

We observed that for the simulated scenario with 7 time points, DynOmics increased sensitivity compared to commonly used methods by at least 8%, while still remaining highly specific (*>* 0.9). With 14 time points, all methods performed similarly, including DynOmics. Pearson correlation which does not take time delays into account performed the worst in terms of sensitivity in all scenarios, demonstrating that ordinary correlation measures are not sufficient to detect associations when trajectories are delayed. A detailed description of the simulation study and the results is provided in the Supporting Material Section A.

### Implementation and computation time

DynOmics is implemented in R and uses the FFT implemented in the function `fft()` from the `stats` R package^49^ for the decomposition of the time series. DynOmics utilises the R package `parallel` to perform calculation on CPUs in parallel where possible. DynOmics’ computation time was tested and compared to DTW4Omics on simulated datasets with seven time points and the Lung Organogenesis study described below with 14 time points. On simulated data with one reference and 100 queries DynOmics required two seconds, while DTW4Omics required 30 seconds. On the Lung Organogenesis study the association of 50 references and 50 queries took DynOmics four seconds compared to 600 seconds for DTW4Omics.

## Case studies

### Lung Organogenesis

#### Description of the study

The study of Dong *et al.*^34^ investigated the dynamic regulation of miRNAs in mouse lung organogenesis by measuring the expression of 516 miRNAs and 45,105 mRNAs on two biological replicates at seven time points (embryo day 12,14,16,18; postnatal day 2,10,30) in lungs (GSE21053, Affymetrix Mouse Genome 430 2.0 Array). The data we analysed were pre-processed in the original study. Subsequently, a linear mixed effect model splines (LMMS) modelling approach developed previously^3^ was used to obtain representative trajectories over 14 equally spaced time points between embryo day 12 and postnatal day 30. In addition to allowing interpolation to even out spacing between time points, LMMS can handle unbalanced designs - when the number of observations per time point is unequal, or if there are missing data. We further filtered the data to retain only miRNAs and mRNAs declared as differentially expressed over time using lmmsDE^3^ (FDR *≤* 0.05). The final dataset analysed with DynOmics included 105 miRNAs and 11,326 mRNAs.

#### Analysis strategy

We compared associations detected between miRNA and mRNA pairs for both raw and LMMS modelled trajectories, using either classical Pearson correlation (on raw and LMMS modelled data) or DynOmics (on LMMS modelled data). MiRNAs are known to be able to target transcription regulators and therefore lead to the indirect expression of many mRNAs downstream.^30^ In this study, however, we focused on direct targets of miRNA, and therefore sought to identify negative correlations between miRNAs and mRNAs, *i.e.*, increased miRNA expression levels associated to a decreased (inhibited) mRNA expression levels, or vice versa. Associations were declared for all miRNA-mRNA pairs whose Pearson correlation coefficient was *<* −0.9. The mRNAs associated to a given miRNA were compared with miRNA targets predicted from sequence similarity from microRNA.org (GoodmirSVRscore, Conserved miRNA, release August 2010),^50^ TargetScan (release 6.2)^51^ and miRDB (Version 5).^52^ We only compared database entries with an exact identifier match to the analysed 105 miRNAs, leaving 14 miRNAs for miRDB, 33 for TargetScan and 86 for microRNA.org. Pathway enrichment analysis was performed using QIAGEN’s Ingenuity^®^ Pathway Analysis (IPA^®^, QIAGEN Redwood City, www.qiagen.com/ingenuity).

### Mammalian Embryonic Development

#### Description of the study

Xie *et al.*^36^ investigated the dynamic expression of human, mouse and bovine transcripts during PED. The expression levels in human (30,283 mRNAs), mouse (19,607 mRNAs) and bovine (13,898 mRNAs) were monitored during six to eight comparable cell stages (oozygote (only bovine), zygote, two-, four-, eight-, 16-cell (bovine only), morula and blastocyte) in two (human, mouse) and three (bovine) embryo replicates (GSE18290, Affymetrix mircoarrays: Mouse Expression 430A Array, Human Genome U133 Plus 2.0, Bovine Genome Array).

#### Analysis strategy

We first converted the cell stages (zygote to blastocyte) into quantitative time points (one to seven) for input into modelling. For each organism, expression trajectories were modelled using LMMS with 14 regularly spaced time points. Human transcripts were taken as references, with reference-query pairs restricted to orthologous sequences with mouse and bovine as specified in the Affymetrix file HG-U133 Plus 2.na26.ortholog.csv. Seven human transcripts did not match any identifier in the orthology file and were removed. A total of 81,966 orthologous transcript pairs were analysed (48,566 mouse, 33,400 bovine), where references and/or queries may have been included in multiple pairs. We applied DynOmics to every orthologous transcript pair to assess delays in expression levels between organisms and declared association when the absolute correlation exceeded 0.9. Pathway enrichment analysis was performed using IPA.

## Results

### Lung Organogenesis

Firstly, focusing on the miRNAs as reference trajectories, we compared the performance of Pearson correlation on the raw and LMMS modelled data. We defined the average agreement as the number of associations identified in common between the two methods divided by the number of associations observed by one method averaged over all miRNAs (Supporting Table S3). We found that modelling representative trajectories using LMMS substantially increased the number of associations, by over 80% compared to raw data. This is likely due to the removal of noise when modelling the trajectories.^3^ We next compared the performance of Pearson correlation with DynOmics for the LMMS modelled data. DynOmics identified on average 18% more associations, indicating that the simple correlation analysis was not sufficient to detect all delays in expression between miRNA and mRNA.

Secondly, we analysed the overlap of these putative miRNA targets with the miRNA targets predicted from sequence similarity. Supporting Tables S4-S7 summarise for each miRNA and each method the number of putative targets and the overlap with the predicted targets from TargetScan, microRNA.org and miRDB. For the raw data, we observed low overlap between predicted and putative targets (ranging from 0 to 0.4% miRDB, 1.8% microRNA.org, and 4.8% TargetScan). The number of overlaps increased for the LMMS modelled data with the majority of miRNA-mRNA pairs changing expression simultaneously (*i.e.*, a delay of 0). However, the percentage of overlap was still small (ranging from 0 to 3% miRDB, 3.5% microRNA.org, and 4.8% TargetScan; Supporting Figure S5).

Finally, we investigated whether the putative delays were of biological relevance for miRNA-mRNA pairs. Three miRNAs in particular, mu-miR-429, mmu-let-7g, and mmu-miR-134, were associated with a large number of negatively delayed mRNAs, represented in Figure 2 1-3. Analysis of these delayed mRNAs using IPA identified for mmu-miR-429 enrichment of the ‘Phospholipase C Signaling’ pathway (*P* = 1.21×10^*−*14^), for mmu-let-7g the ‘Axonal Guidance Signaling’ pathway (*P* = 4.0 × 10^*−*11^), and for mmu-miR-134 the ‘Mitotic Roles of Polo-Like-Kinase’ pathway (*P* = 1.29×10^*−*8^). These pathways have been described as being involved in either embryonic or lung development. Phospholipase C was associated with fetal lung cell proliferation in rats^53^ and plays an important role in organogenesis and embryonic development.^54^ Some axonal guidance molecules like netrins have been suspected to play a role in lung branching,^55^ while EphrinB2 or semaphorin 3C were found to be involved in alveolar growth and development.^56, 57^ Finally, polo-like-kinases (PLKs) are highly conserved in mammals and are important for early embryonic development.^58^ PLKs are known to regulate cell cycle progression but little is known about their role in lung development. However, over-expression of PLKs has been associated with malignancy and poor prognosis in lung cancer, and PLKs are therefore a target for therapy.^59^

**Figure 2.**
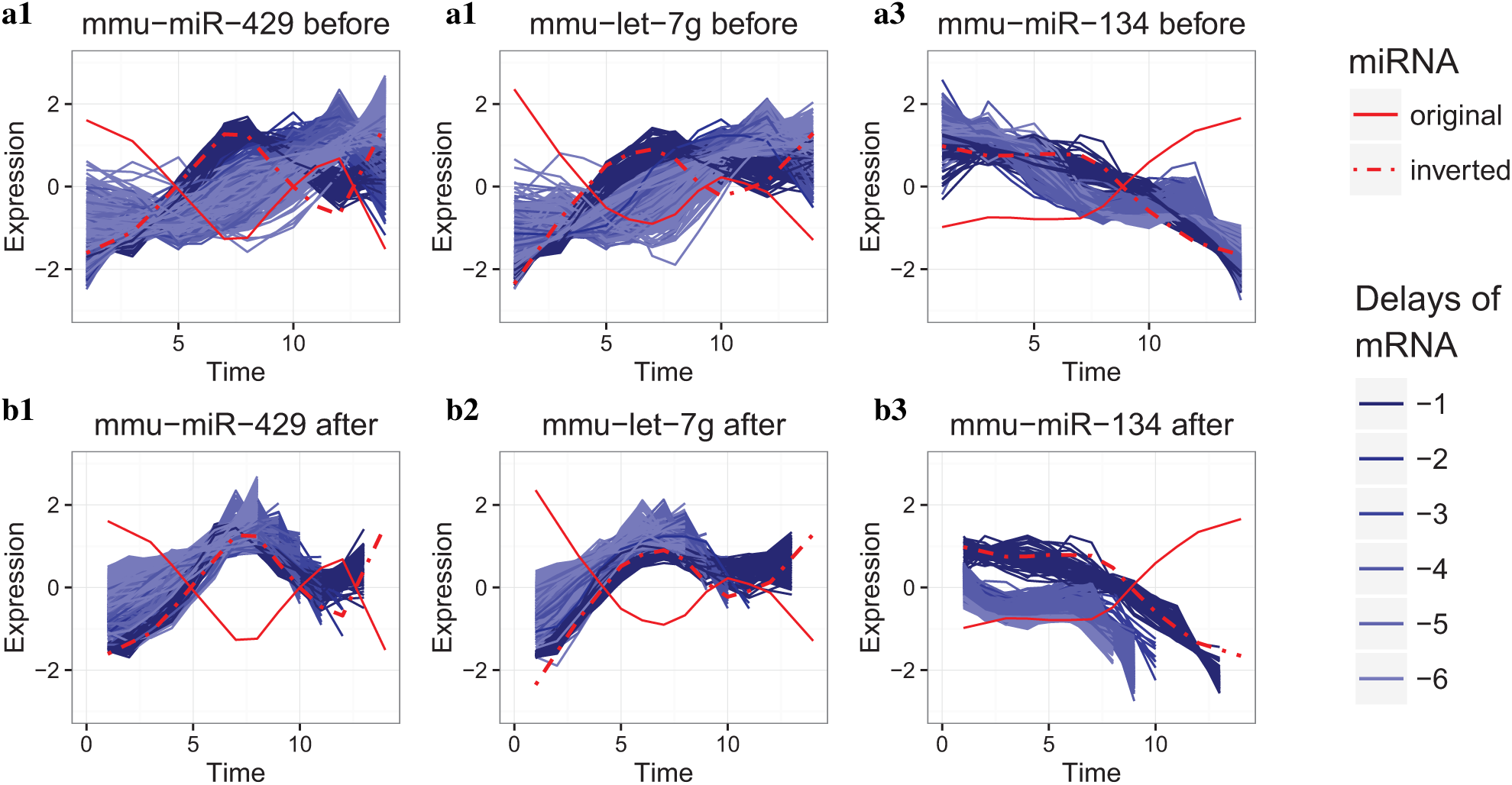
MiRNA and mRNA expression associations in Lung Organogenesis study. Scaled LMMS modelled expression levels (y-axis) are depicted over time in 14 equally spaced time units from embryo day 12 to postnatal day 30 (x-axis) for the miRNAs mmu-miR-429, mmu-let-7g, and mmu-miR-134 (red lines). Solid lines depict actual scaled expression levels, while dashed lines depict inverted scaled expression levels to account for the negative correlation with mRNA. Modelled expression levels of the mRNAs identified as associated with each miRNA using DynOmics are displayed (DynOmics correlation *<* −0.9, delay *<* 0) **a**) before and **b**) after shifting the trajectories using the DynOmics estimated delay. The blue color gradient reflects the amount of delay.

### Mammalian Pre-implantation Embryonic Development

We applied DynOmics to identify delays in orthologous transcript expression of mouse and bovine relative to human during PED. For an absolute correlation threshold of 0.9, we identified 32,329 (67%) orthologous pairs as being associated between human and mouse, and 26,769 (80%) between human and bovine, summarised in Table 1 with respect to the different types of delay. Of the transcripts displaying association, we observed that the majority of the mouse (56%) and bovine (67%) transcripts were not delayed compared to the orthologous human transcripts. Interestingly, 20% of mouse transcripts (compared to 10% in bovine) changed expression prior to the human orthologous transcript. This could reflect timing differences in the zygote genome activation of mouse PED at the gene expression level.^36, 37^

**Table 1.** Orthologous transcripts identified as associated by DynOmics. Number (percentage) of mouse and bovine transcripts identified as associated with orthologous human transcripts at an absolute correlation threshold of 0.9. The number of associations are divided according to different types of delay, indicating whether changes in expression levels of the mouse and bovine transcripts occurred prior to (delay*>* 0), simultaneously to (delay= 0), or after (delay*<* 0) expression changes of the orthologous human transcript.

| Delay | Mouse vs Human (%) | Bovine vs Human (%) |
| --- | --- | --- |
| > 0 | 6,582 (20) | 2,766 (10) |
| 0 | 18,065 (56) | 17,906 (67) |
| < 0 | 7,682 (24) | 6,097 (23) |
| Total | 32,329 | 26,769 |

Pathway analysis using IPA was performed on the human orthologs for the three types of delay (negative, no delay and positive) relative to mouse or bovine orthologs. Table 2 lists the top three canonical pathways identified as enriched for each type of delay and organism. The majority of trajectories whose expression levels changed in mouse prior to human were involved in EIF2 Signaling (*P* = 7.94 × 10^*−*18^), mTOR Signaling (*P* = 5.64 × 10^*−*12^) and regulation of eIF4 and p70S6K Signaling (*P* = 5.72×10^*−*11^). EIF2 Signaling and eIF4 and p70S6K Signaling play an important role in translation regulation and mTOR Signaling is an important pathway in embryonic development.^60^ These same pathways were also highlighted in a recent study using RNA-Sequencing technologies^61^ on human during early embryonic development (4-cell, 8-cell, morula, and blastocyte stages). EIF2 Signaling (*P* = 1.75 × 10^*−*25^) and the regulation of eIF4 (*P* = 3.48 × 10^*−*0.9^) were also found to be enriched in bovine; however, the genes involved in these pathways changed expression after human expression changes.

**Table 2.** IPA enrichment analysis of human orthologs for three types of delay relative to mouse/bovine transcripts. The top three IPA enriched pathways are listed. Associated transcripts were analysed separately with respect to the delay: positive (negative) delay indicates that the mouse or bovine ortholog’s expression changes occurred prior to (after) the human expression changes. No delay indicates that all expression changes occurred simultaneously.

| Delay compared to human | Organism | Pathway | P value |
| --- | --- | --- | --- |
| > 0 | Mouse | EIF2 Signaling | $7.94 \times 10^{-18}$ |
| | | mTOR Signaling | $5.64 \times 10^{-12}$ |
| | | Regulation of eIF4 and p70S6K Signaling | $5.72 \times 10^{-11}$ |
| | Bovine | Protein Ubiquitination Pathway | $6.28 \times 10^{-09}$ |
| | | Amyloid Processing | $4.36 \times 10^{-08}$ |
| | | Glucocorticoid Receptor Signaling | $1.03 \times 10^{-06}$ |
| 0 | Mouse | Role of Macrophages, Fibroblasts and Endothelial Cells in Rheumatoid Arthritis | $1.84 \times 10^{-31}$ |
| | | Role of Osteoblasts, Osteoclasts and Chondrocytes in Rheumatoid Arthritis | $6.88 \times 10^{-27}$ |
| | | Axonal Guidance Signaling | $1.29 \times 10^{-22}$ |
| | Bovine | Protein Kinase A Signaling | $7.11 \times 10^{-16}$ |
| | | Thrombin Signaling | $1.44 \times 10^{-15}$ |
| | | Acute Phase Response Signaling | $4.51 \times 10^{-15}$ |
| < 0 | Mouse | Ephrin Receptor Signaling | $8.39 \times 10^{-13}$ |
| | | Molecular Mechanism of Cancer | $5.26 \times 10^{-12}$ |
| | | B Cell Receptor Signaling | $1.54 \times 10^{-10}$ |
| | Bovine | EIF2 Signaling | $1.75 \times 10^{-25}$ |
| | | Regulation of eIF4 | $3.48 \times 10^{-09}$ |
| | | Protein Ubiquitination Pathway | $5.11 \times 10^{-09}$ |
| > 0, 0, < 0 | Mouse, Bovine | EIF2 Signaling | $1.59 \times 10^{-17}$ |
| | | Regulation of eIF4 | $5.25 \times 10^{-10}$ |
| | | Acetyl-CoA Biosynthesis I (Pyruvate Dehydrogenase Complex) | $8.71 \times 10^{-06}$ |

As an illustrative example we display the trajectories of the orthologous transcripts involved in EIF2 Signaling in human and mouse with respect to the type of delay (Figure 3).

**Figure 3.**
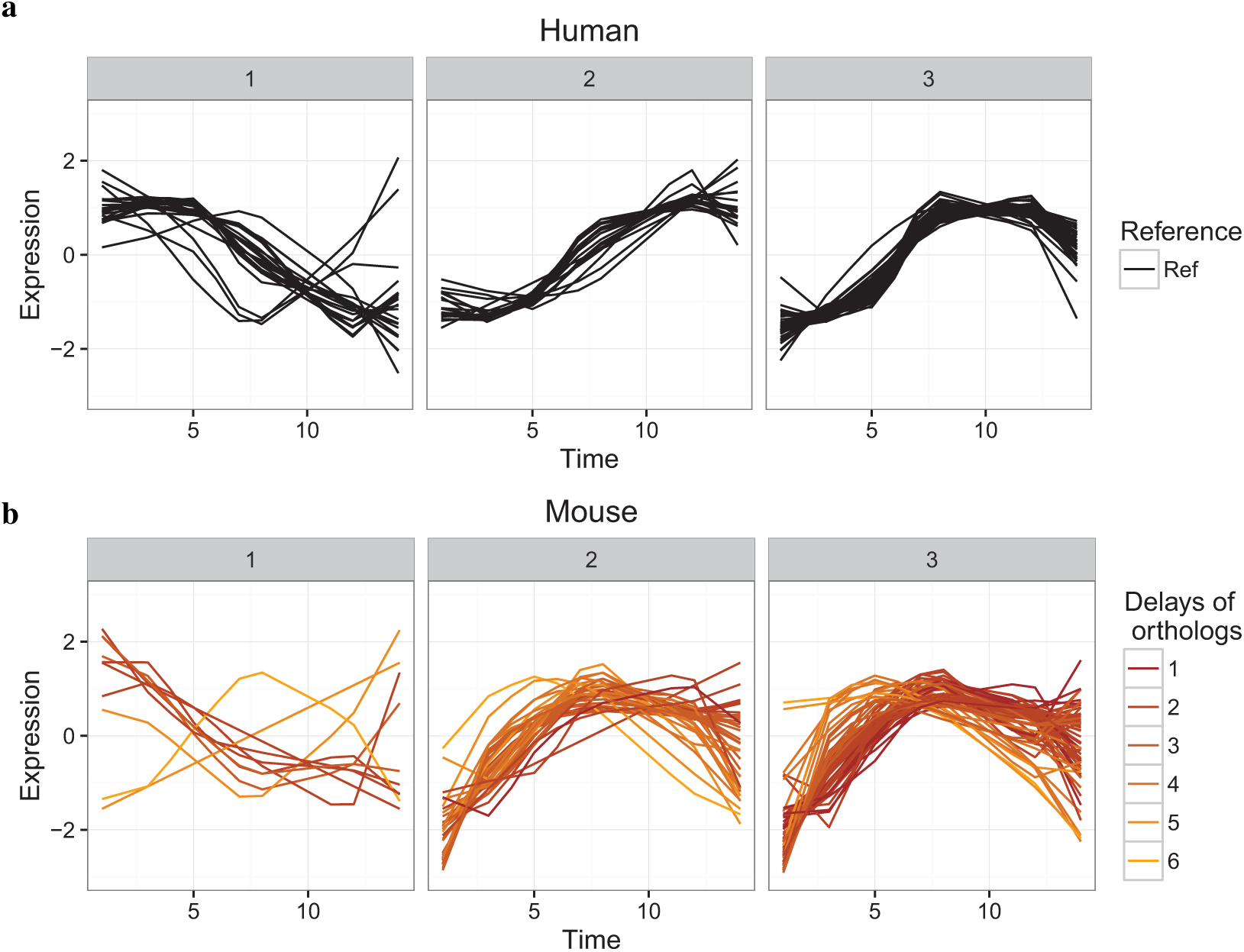
EIF2 Signaling. Modelled transcripts expression levels (scaled for each time point for visual purposes, y-axis) with respect to time (x-axis) involved in EIF2 Signaling **a**) in human with **b**) their orthologs in mouse (DynOmics correlation *>* 0.9, delay *>* 0). Hierarchical clustering was performed on the human transcripts to extract three main expression patterns in EIF2 Signaling (**a**; 1-3). The three main patterns of expression in humans **a)** were visualised in separate plots (1-3). The mouse expression profiles in **b**) were separated by the classification of their human orthologs (1-3) and were coloured according to the DynOmics estimates of delay.

We also performed enrichment analyses for human orthologs for all transcripts identified as associated, across all three types of delay. We highlight the conserved process of Acetyl-CoA Biosynthesis I, since it has not occurred in the enrichment looking at the delayed orthologs individually. Acetyl-CoA levels were found to play a role in the acetylation of proteins and may play a role in regulation of embryogenesis.^62^ Using DynOmics, we identified different response dynamics across organisms for four out of six transcripts (dihydrolipoamide branched chain transacylase (DBT), dihydrolipoamide s-acetyltransferase (DLAT), dihydrolipoamide dehydrogenase (DLD) and pyruvate dehydrogenase (Lipoamide) beta (PDHB)) that are conserved and involved in this process (Table 3).

**Table 3.**
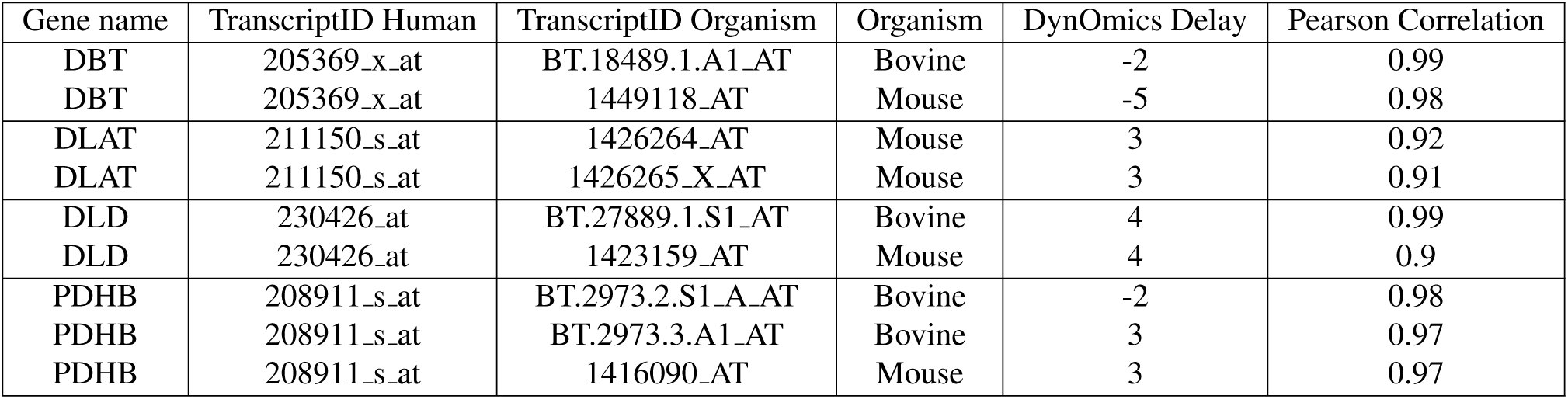
Acetyl-CoA Biosynthesis I orthologous transcripts. Orthologous transcripts identified as associated by DynOmics and involved in the Acetyl-CoA Biosynthesis I pathway. Gene names, transcript IDs in human, bovine and mouse are indicated, as well as the estimated DynOmics delay and the Pearson correlation between the reference trajectory and the query trajectory after shifting based on the DynOmics delay estimate.

## Discussion

To date, very few methods have been developed to integrate time course ‘omics’ data that are robust to delays in expression between co-expressed molecules. The integration task is particularly challenging as the data are often characterised by a high level of noise and measured on a small number of time points. Our algorithm DynOmics addresses these challenges by modelling time course trajectories, identifying delays and re-aligning trajectories to determine the degree of mutual dependency between reference and query trajectories.

Modelling time course trajectories is an important step in this process, as most methods developed to integrate time course data, such as DTW4Omics^24^ and HMMs^25^ require as input only a single value per time point. In this study we used a data-driven modelling approach based on linear mixed model splines^3^ to summarise the time course data appropriately, to reduce noise, and to interpolate additional time points within the time course. We found that while the modelling step may remove some associations between reference and query trajectories, *e.g.*, in the Lung Organogenesis case study, it ultimately increased the number of findings by considerably reducing the amount of noise in the data. In addition, modelling each trajectory as a noisy function of time allows integration of datasets with different time intervals or numbers of time points, as we demonstrated in the mammalian embryonic development case study.

The selection of an appropriate threshold to define associations between co-expression trajectories is not trivial, and depends on the characteristics of the data themselves. For our analyses, we specified a correlation threshold of 0.9, as we were only interested in highly concordant expression trajectories.

The role of miRNAs as gene expression regulators is an exciting new subject of study, as it is estimated that they control one-third of the expression of the human genome.^63^ Moreover, since miRNAs appear to be the master switch in biological processes, they are the target of future therapeutic development.^31, 32^ In the Lung Organogenesis study, the miRNA-mRNA associations that we identified with DynOmics largely did not agree with the predictions from the databases TargetScan, microRNA.org and miRDB. One possible reason for the general lack of agreement could be that the predicted targets were not expressed in the experiment, or were not targeted at all. Those mRNA that did agree represented subtle delays between miRNA and mRNA trajectories that may indicate high sequence affinities. The other associations, which included larger delays that were not identified by standard correlation analysis,^34^ may not be as similar in sequence and hence were not predicted as miRNA targets in the databases.^50–52^ Indeed, our results suggest that sequence information alone may not suffice to determine whether miRNAs are expressed and regulate specific mRNA under certain conditions. Alternately, the large number of miRNA-mRNA associations identified by DynOmics may represent mRNAs which are indirect targets of miRNAs. Determining whether these mRNAs are truly direct targets of miRNAs will require further experimental validation, but the enrichment analysis showed that the mRNAs were involved in meaningful biological processes related to Lung Organogenesis, *e.g.*, lung cell proliferation, lung branching and alveolar development. Thus, in this context, DynOmics has the potential to identify novel targets of miRNAs to aid in therapeutic development. Our study emphasises the importance of jointly studying miRNA and mRNA expression to understand the mechanisms of miRNA regulation.

Model organisms present a simpler and more convenient alternative to directly study disease in humans. In the mammalian pre-implantation embryonic development study, we showed that DynOmics could identify delayed conserved expression between different organisms. This is a challenging task, as timing differences of expression changes can occur both in metabolic processes and across organs for different organisms.^37^ By correcting for these timing differences, DynOmics can therefore help to infer gene functions across organisms, and thereby integrate information in whole biological processes. Such integration may in turn identify which organisms provide suitable models for human disease and drug discovery due to the conservation in processes.^64^

Currently, DynOmics has been used to identify associations between datasets of moderate size (~100 references and ~10,000 queries). The computational time would increase for large data sets (~10,000 references and queries). One solution could be to cluster profiles prior to applying DynOmics, to identify specific patterns of interest over time as queries and/or references. As the algorithm is based on independent pairwise comparisons, parallel computing could also be used to decrease the computational burden. Alternatively, as shown in the Lung Organogenesis study, the DynOmics analysis can be performed on a smaller number of queries selected based on prior knowledge or biological assumptions.

## Conclusion

Delays in molecular expression are an acknowledged and important phenomenon in many areas of biology. Here we demonstrated the need for and value of methods that are robust to delays, by showcasing the benefit of accurate delay estimates to interpret response dynamics and identify conserved molecular mechanisms. DynOmics overcomes the challenge of integrating data with timing differences of expression changes and therefore presents an effective tool to study time-sensitive molecular expression. The integration of multiple time course ‘omics’ data is becoming necessary in order to understand a biological system’s formation, actions and regulation with high confidence. Our algorithm DynOmics provides a unique opportunity to study molecular interactions between multiple functional levels of a single system or multiple organisms. DynOmics is implemented in the open source programming language R and is freely available via bitbucket.

## Acknowledgements

This work was supported by the Wound Management Innovation CRC (established and supported under the Australian Government’s Cooperative Research Centres Program) [JS], the Australian Cancer Research Foundation (ACRF) for the Diamantina Individualised Oncology Care Centre and National Health and Medical Research Council Career Development (NHMRC) fellowship [APP1087415 to KALC] and the Australian Research Council [DE120101127 to BEH].

## Contributions

JS developed the methodologies, implemented the approaches, performed the statistical analyses and wrote the manuscript. KALC and BEH helped writing the manuscript. All authors participated in the design of the study and reviewed the manuscript.

